# PREP-aring is worth it: Success of the Case Western Reserve University Postbaccalaureate Research Education Program and its Scholars

**DOI:** 10.64898/2025.12.30.697004

**Authors:** Dana C. Crawford, Esteban Vazquez-Hidalgo, Hua Lou

## Abstract

The National Institute of General Medical Sciences (NIGMS) Postbaccalaureate Research Education Program (PREP) was a research-intense, one-year training program for recent college graduates from experiential backgrounds uncommon in science who intended to matriculate with a PhD program in preparation for a career in biomedical research. Case Western Reserve University had an NIGMS-supported PREP (CasePREP) from 2007 until 2025, the year NIGMS terminated the program and early expired the PREP funding mechanism. We report here the extent that CasePREP, with its 108 Scholars, met program goals as well as the economic and scientific impact NIGMS-supported PREP has had for a variety of stakeholders. CasePREP was a great success as measured by its major outcomes: matriculation into a PhD or MD/PhD program (83%) and completion of degree (51 Scholars to date). CasePREP Scholars have persisted in science and have made substantial contributions to the scientific workforce and enterprise. PREPs in general provided a crucial bridge for research talents who had little prior research opportunities to realize their potential and career goals. NIGMS termination of PREP support has negative economic consequences and endangers an important pathway to a career in science for Americans who would not otherwise have similar opportunities.

## Introduction

The Postbaccalaureate Research Education Program (PREP) was established in 2000 by the National Institute of General Medical Sciences (NIGMS) at the National Institutes of Health with the funding announcement PAR-00-139 and first awards in 2001 [1,2].

The intention and goals of the program targeted well-documented scientific workforce challenges [3] by providing research and professional development opportunities to recent college graduates from experiential backgrounds not common in science who were interested in pursuing a doctoral degree in biomedical research fields of study. The program was initially intended for specific US demographic groups but had since expanded to include uncommon backgrounds based on geography (e.g., rural), economic resources, disabilities, and other factors or situations that negatively impacted academic performance and/or limited research opportunities necessary for successful matriculation into a PhD-granting program in biomedical research.

Case Western Reserve University (CWRU) School of Medicine was awarded PREP in 2006 under PAR-03-140 and has been ongoing through 2025 under PAR-17-051. CWRU (“Case”) PREP has completed 18 cohorts of “Scholars” and is one of the longest running NIGMS-support PREPs in the United States. With almost 20 years of data for 108 CasePREP Scholars, we describe here the program outcomes as well as the economic and scientific impact CasePREP has had on the state of Ohio and the international research community.

## Methods and Materials

An overview of CasePREP as a training program is given in **Appendix A** available in **Supplementary Materials**. Outcomes data were collected from Scholars 1) during the application process, 2) during CasePREP, and 3) after CasePREP. Required application data included basic demographic data, permanent and temporary contact information, self-reported eligibility for CasePREP, undergraduate institution, grade point average, degree, personal statement, statement of interest, and letters of recommendation.

Optional was self-reported standardized test scores. Data collected during CasePREP included pre- and post-PREP surveys, courses taken during PREP, grades, seminars attended, laboratory rotations, research summaries, oral and poster presentations given, conferences attended, awards, publications, applications to graduate programs and decisions, and immediate post-PREP position taken. Surveys were self-administered; other data were self-reported in a custom database. Post-PREP data were collected through personal contact, public-facing websites and social media, PubMed, and NIH Reporter. Where indicated, large language models were used for text processing and exploratory categorization; all interpretations and conclusions were determined by the authors.

Economic impact estimates of CasePREP are given in **Appendix B**. Carnegie Classifications of Institutions of Higher Education were used to characterize Scholars’ final degree granting undergraduate institutions and institutions enrolled by Scholars for PhD or MD/PhD programs using their “institutional characteristics” and “student access and earnings” columns. Salaries were estimated for CasePREP Scholar alumni who had a post-terminal degree position as of December 2024. Actual salaries were collected from public sources, and when not available, estimates were collected from public sources (Glassdoor, Salary.com) based on the Scholar’s employer and position. When lower and upper salaries were available, the average of the two was used to represent the Scholar’s estimated salary.

CasePREP Scholar publications and associated PubMed IDs were collected manually using Scholar names and PubMed. Publication titles associated with the PubMed IDs were extracted from PubMed and parsed using CRAN packages ‘easyPubMed’ and ‘wordcloud’ in R scripts generated using Anthropic’s Claude Sonnet 4.5 and subsequently reviewed and modified by the authors. Data were plotted using ggplot2 in R 4.5.1. Citation and publication influence metrics were based on iCite queries [4] using CasePREP Scholar-associated PubMed IDs as inputs. iCite also provided “cited by” PubMed IDs, which were used to calculate first and second-generation citation metrics through iCite and requested Claude Sonnet 4.5 analyses.

The CWRU Institutional Review Board (IRB) determined CasePREP (STUDY20240922) to be non-human subjects research.

## Results

CasePREP appointed 108 Scholars between the 2007-2008 and 2024-2025 academic cycles. Approximately one-third (33%) were male, and all were US citizens or permanent US residents not more than 36 months from when their bachelor’s degree was conferred. At the time of application, Scholars resided in 25 states, Puerto Rico, and the US Virgin Islands, with the majority residing in Puerto Rico (38.89%), Ohio (10.19%), California (9.26%), and New York (5.56%; **Figure 1**).

**Figure 1.**
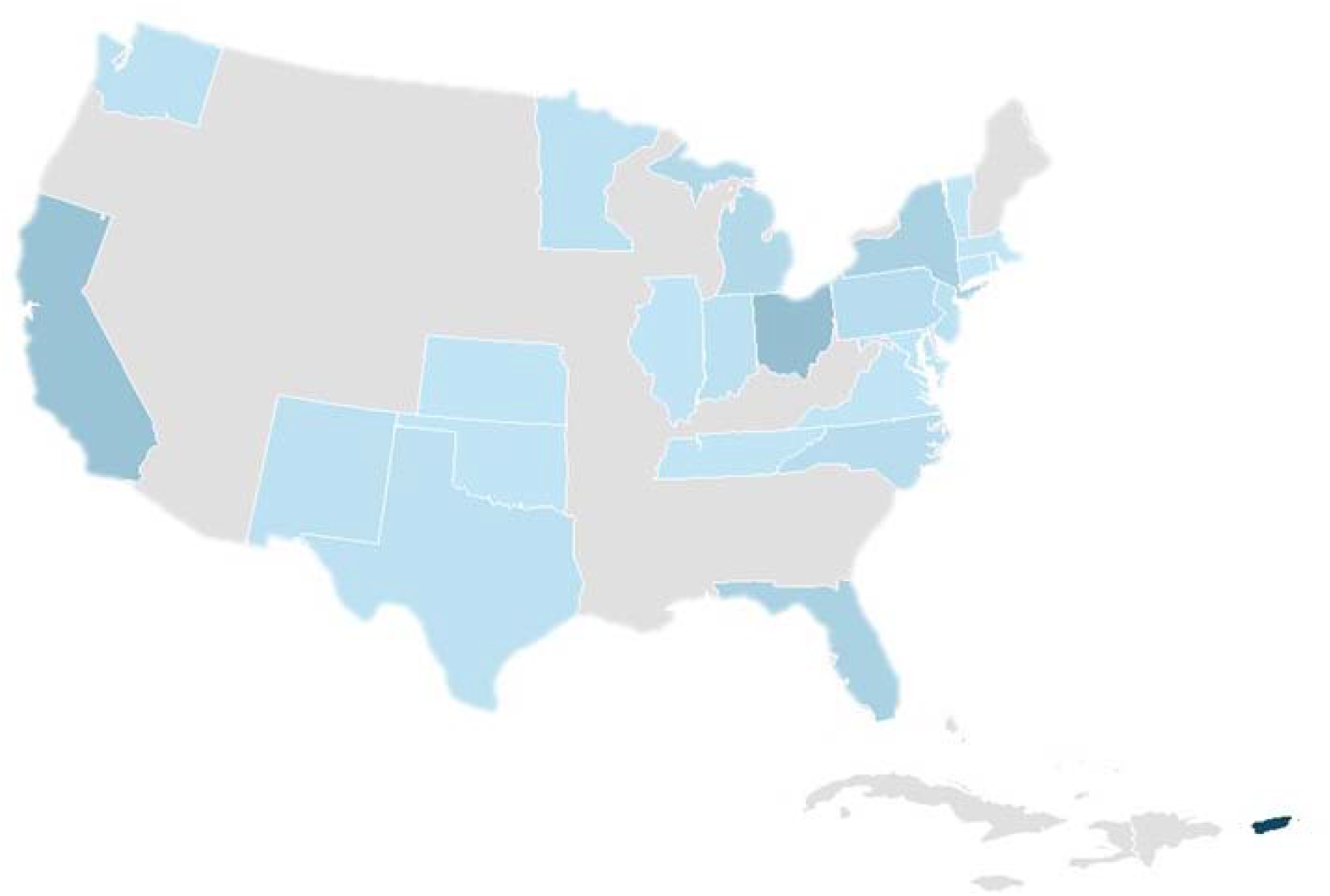
Home state of CasePREP Scholars. US states and territories colored blue represent at least one CasePREP Scholar’s home state at the time of application to the program. The shade of blue represents proportion of Scholars residing in the state from least (light blue) to most (dark blue). States shaded grey do not have residing Scholars. Not shown: Alaska (grey), Hawaii (grey), and the US Virgin Islands (light blue).

CasePREP Scholars earned four-year Baccalaureate degrees from 58 US institutions in 21 states, Puerto Rico, and the US Virgin Islands. According to the Carnegie Classifications of Institutions of Higher Education, the majority of the institutions are classified as “mixed Undergraduate/Graduate-Doctorate” (43.52%) and “mixed baccalaureate” (17.59%) (**Appendix C**). Of the 58 institutions, most were classified as having research activities: “Research 1: Very High Spending and Doctorate Production” (30.56%), “Research 2: High Spending and Doctorate Production” (36.11%); and institutions spending at least 2.5 million on research & development on average in a single year but not classified as Research 1 or 2 (6.48%). A quarter (26.85%) of CasePREP Scholars’ baccalaureate institutions do not have substantial research activity.

Almost all (98.15%) CasePREP Scholars completed the post-baccalaureate program. Of those who completed CasePREP, the majority (83%) matriculated into a PhD or MD/PhD program within one to two years of completing the post-baccalaureate program. CasePREP Scholars matriculated with PhD and MD/PhD programs across 33 US institutions, including CWRU School of Medicine (30.7%) and the University of Michigan Medical School (10.2%). Of the 88 who entered a PhD or MD/PhD program, at least 18 (20.45%) received independent, external fellowships or supplements for their training, including 12 NIH National Research Service Award (NRSA) F31s; one NIH NRSA F30; two National Science Foundation (NSF) Graduate Research Fellowship Program (GRFP) fellowships; and three Howard Hughes Medical Institute (HHMI) Gilliam Fellowships. Another two trainees were awarded grants to transition from predoctoral studies to post-doctoral studies (NIH F99/K00). Three CasePREP alumni with PhDs are now Burroughs Wellcome Fund (BWF) Postdoctoral Diversity Enrichment Program (PDEP) awardees (with one also designated a Revson Scholar).

### CasePREP Scholar Contributions to Science

Of the 108 CasePREP Scholars, 75 (69.44%) are co-authors on a manuscript published in a peer-reviewed journal indexed in PubMed. As of September 2025, there are 407 unique PubMed IDs associated with CasePREP Scholars; of these, 15 PubMed IDs are associated with more than one CasePREP Scholar reflecting CWRU mentors and research groups who trained both CasePREP Scholars and alumni as graduate students. Based on manuscript titles, the most common research topics published by CasePREP Scholars includes cancer research (n=57 mentions of “cancer”). Other areas of research and fields of study are also represented, including immunology research (n=26 mentions of “immune” or “immunity”) and COVID-19 research (n=17 mentions of “covid”); research using model organisms (n=18 mentions of “mice” or “rats”); and nanotechnology research (n=22 mentions of “nanoparticles(s)”) (**Appendix D**). The number of publications per year peaked in 2021 with 58, possibly reflecting the consequences of the COVID-19 pandemic stay-at-home mandates in 2020 that reduced laboratory hours and increased work-at-home hours conducive to writing. In general, the number of publications has increased over the course of CasePREP (**Figure 2**), both reflective of the increasing cumulative number of Scholars in the long-running CWRU post-baccalaureate program but also the retention of CasePREP alumni in graduate school and in the scientific workforce afterward.

**Figure 2.**
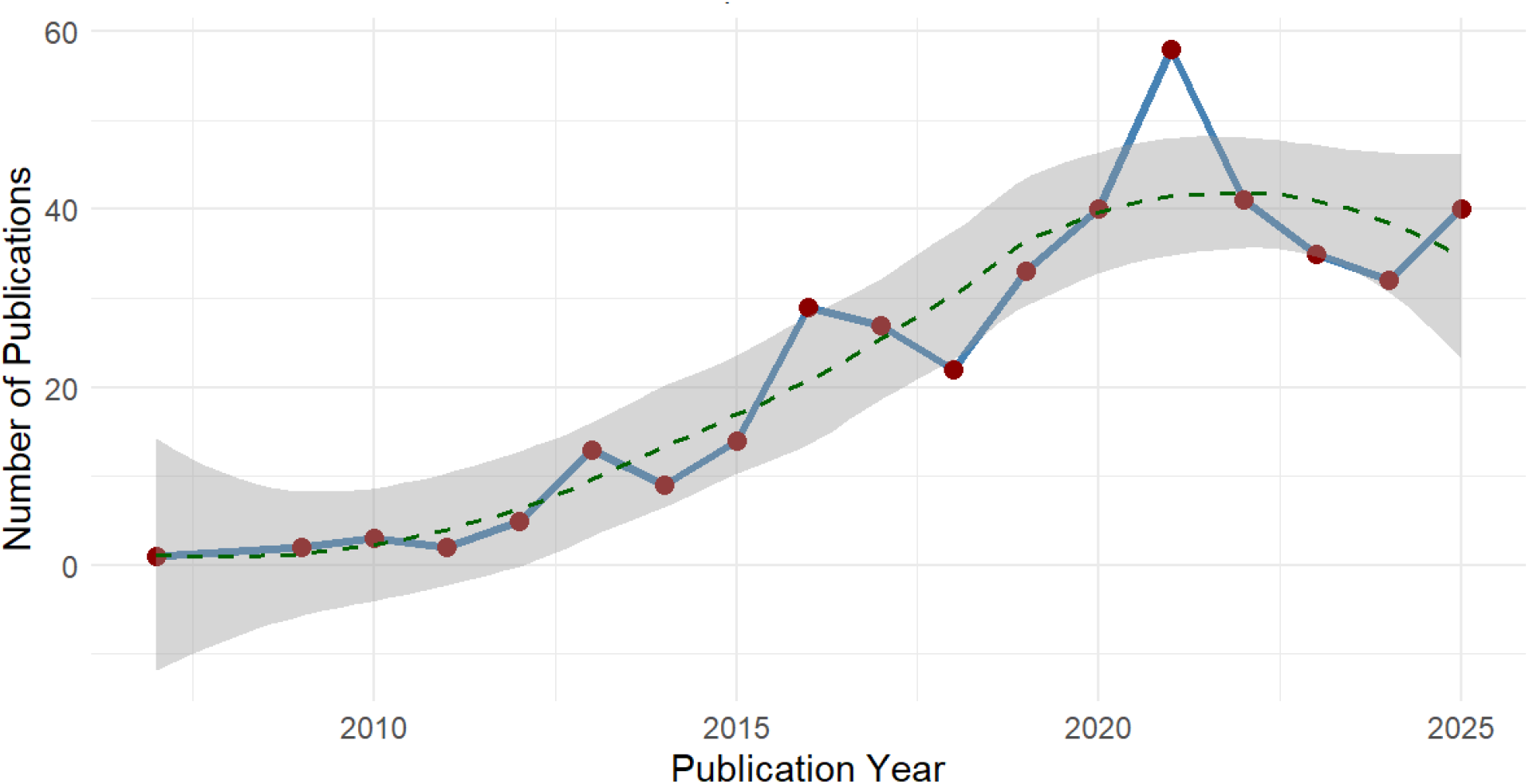
CasePREP Scholar scientific publications over time. Plotted are the number of publications published by CasePREP Scholars (y-axis) by publication year (x-axis). Publications were identified through PubMed by CasePREP Scholar alumni names and include publications based on research prior to CasePREP, during CasePREP, and after CasePREP. PubMed indexed bioRxiv and medRxiv manuscripts are not included in this plot. As of the end of September 2025, 75/108 CasePREP Scholars had at least one PubMed indexed manuscript published in a peer-reviewed journal. The plot was generated by ggplot in R using geom_smooth (method = “loess”, se = TRUE), which computes the regression line (dashed line) and 95% confidence interval (gray) for the total number of publications per year (red dots) over time.

To characterize the impact of CasePREP Scholar research, we queried iCite [4] for number of citations, relative citation ratios, and PubMed IDs of citing publications. Of the 407 CasePREP Scholar publications, 406 were available in iCite. As of September 2025, the Scholars’ published work has garnered 14,261 total citations, with an average 35.04 citations per publication (standard error of the mean = 3.19). The Scholars’ collective body of work has a weighted citation ratio of 742.59. The most highly cited Scholar-associated publication to date [5] has a relative citation ratio of 24.36, indicating this paper has received 24 times the citations per compared with the median NIH-funded publication. The most highly cited Scholar-associated publication is the most influential CasePREP Scholar-associated publication in a cascade of two generations: it is directly linked to 806 citations (1^st^ generation), and those citations are linked to 43,343 citations (2^nd^ generation). Collectively, the 406 CasePREP Scholar-associated publications are associated with 240,367 2^nd^ generation citations.

### Return of Investment

Over the course of CasePREP, CWRU was awarded a total of $5,623,773 USD ($5,215,025 USD direct costs and $408,708 USD indirect costs) from NIH spanning the parent grant, three renewals, and two supplements. The average NIH investment for 108 CasePREP Scholars on a per Scholar basis was $52,072 USD. Major measures of CasePREP success include the proportion of Scholars who enter graduate programs 1) remain in the programs and 2) complete the programs with a PhD or MD/PhD. As of March 2026, 88 of the 106 (83.02%) Scholars who completed CasePREP entered a PhD or MD/PhD program. One recent CasePREP Scholar deferred enrollment to a PhD program and is expected to matriculate the 2026-2027 academic cycle. Of the Scholars who entered PhD or MD/PhD programs, the majority (85.23%) are either current trainees (n=23) or have earned a PhD (n=46), MS/PhD (n=2), MBA/PhD (n=1), or an MD/PhD (n=2) as their terminal degree. The average time-to-degree for the 51 CasePREP Scholars was 6.24 years: 6.12 (standard deviation = 1.26; median = 6) years for PhDs and 7.40 (standard deviation = 2.30; median = 8) years for dual degrees. Return of investment estimates indicate that NIH costs per CasePREP Scholar who enter graduate school save NIH investments lost to attrition when PhD program attrition rates are high (**Appendix B**).

### Social Mobility and Earning Potential

To characterize realized earning potential, we first considered all CasePREP Scholars who earned a PhD and had a known salaried position as of December 2024 (n=40; **Figure 3**). Of these CasePREP alumni, half held positions in industry and half in academics with an average 4.30 and 3.55 years since completion of their PhD, respectively. Among those with industry positions, the average estimated salary in 2024 was $152,028 USD (standard deviation = $52,403 USD; median = $157,484 USD), ranging from an estimated $74,705 to $238,500 USD. Compared with CasePREP alumni with PhDs in industry, those in academia had a lower estimated average 2024 salary of $74,174 USD (standard deviation = $17,465 USD; median = $71,880 USD), consistent with instructor or junior faculty positions and post-doctoral training.

**Figure 3.**
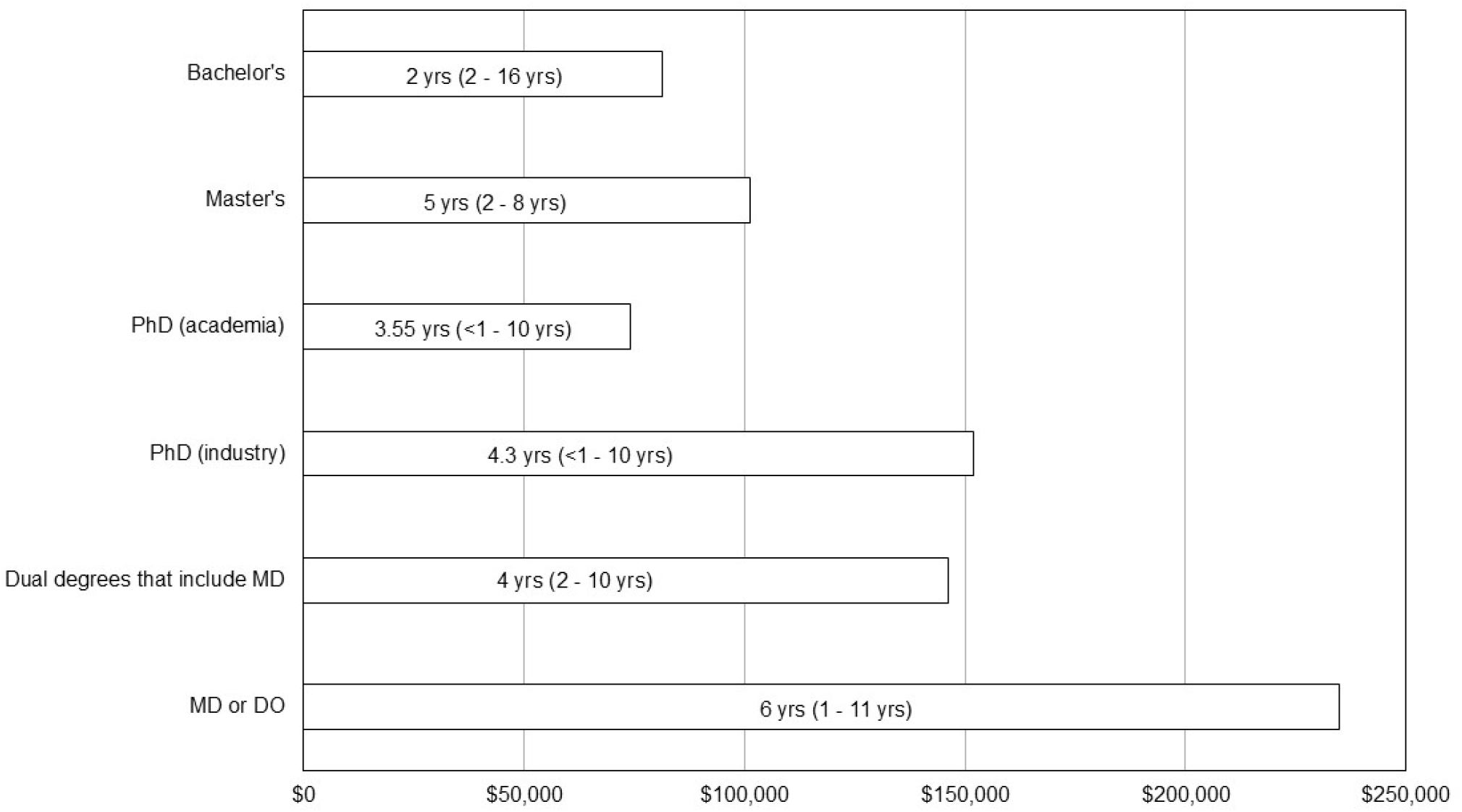
Estimated 2024 annual salaries by terminal degree. We estimated 2024 annual salaries for 63 CasePREP Scholars who had a known salaried position as of the end of 2024. We estimated salaries based on position descriptions posted on LinkedIn and other social media and salary estimated from GlassDoor.com and similar resources (see Methods and Materials). Average salaries are plotted (x-axis) by terminal degree (y-axis), which included Bachelor’s degrees (n=4), Master’s degrees (n=6), PhDs with positions in academia (n=20), PhDs with positions in industry (n=20), dual degree MDs such as MD/PhD, MD/MPH, and MD/MS/JD (n=5), and MDs or DOs (n=6). Not shown are one MBA and one PharmD/MS. Shown in the middle of each salary average is the average number of years since terminal degree (or for Bachelor’s degree, since CasePREP) and the range.

In addition to working CasePREP alumni who earned PhDs, 19 CasePREP Scholars earned other terminal degrees and held salaried positions as of December 2024. These working alumni included those who earned 1) a Master’s (n=6) degree and held positions in industry, 2) a Doctor of Medicine (MD) or Doctor of Osteopathic Medicine (OD) degree (n=6), 3) a dual medical (MD/PhD, MD/JD/MS, or MD/MPH) degree (n=5), 4) a dual PharmD/MS degree, and 5) an MBA. An additional four CasePREP Scholars were known to have entered the workforce with a Bachelor’s degree. The average estimated salary for CasePREP alumni working with a Bachelor’s degree (estimated average of $81,295 USD) is similar to the average 2024 earnings reported by the US Bureau of Labor Statistics for all working Bachelor’s degree holders ≥25 years of age ($80,236 USD) [6]. The salary averages for other terminal degrees are higher than average salary for Bachelor’s degree holders, but these differences are not statistically significant in pairwise comparisons (p>0.05; t-tests) reflecting the wide range of salaries within each terminal degree group and the relatively small sample sizes.

## Discussion

CasePREP, with 108 Scholars and $5.6 million in federal funding over the course of nearly 20 years, has made a substantial impact on the local Cleveland, Ohio economy and the international scientific community. CasePREP alumni include 46 PhDs; 2 MS, PhD; 1 MBA, PhD; 2 MD/PhDs; 6 MDs; 1 DO; 1 MS, JD, MD; 1 MPH, MD; 1 MS, PharmD; 1 MBA; and 9 Master’s (**Figure 4**). More than two dozen alumni are still in training. Post-PREP follow-up indicates that the majority of Scholars, regardless of their terminal degree, entered and have remained in the scientific workforce. For many Scholars, CasePREP made an impactful contribution to their earning potential. Of the Scholars who have published, their collective body of work has been highly impactful reflected in the number of direct and cascading citations in the literature. Collectively these data serve as a direct rejection of the assertion that this program and its Scholars “do nothing to expand our knowledge of living systems, provide low returns on investment, and ultimately do not enhance health, lengthen life or reduce illness.” [7]

**Figure 4.**
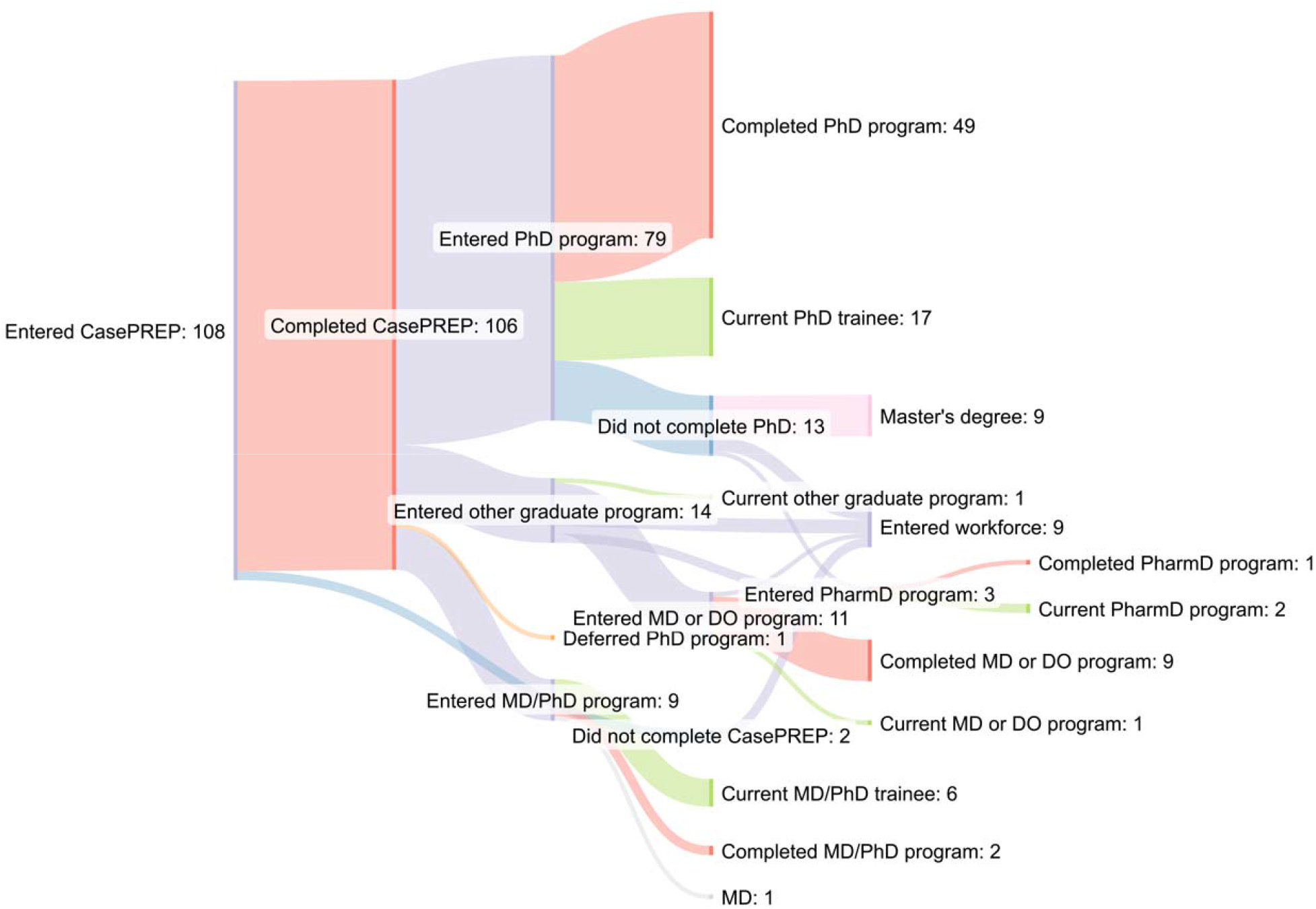
CasePREP outcomes for all Scholars. CasePREP outcomes, as of March - 2026, are shown for all Scholars. Data are visualized as a flow diagram using SanKeyMATIC.

The data presented here were collected primarily for program evaluation purposes; consequently, there are several limitations. First and foremost, CasePREP alumni were not directly surveyed for information such as awards or fellowships, dates of thesis defense, or current salaries. These data were primarily scraped from public-facing LinkedIn and websites or identified in NIH Reporter and NSF GFRP Fellowship database queries. When not available, data were inferred or extrapolated based on reasonable assumptions as detailed in Methods and Materials. While care was taken to provide reasonable point estimates, there may be sizable differences between the estimated and the actual. Also, data available in LinkedIn may not be up-to-date or as granular as desired. For example, the CasePREP average time-to-PhD of 6.12 years is likely closer to the national average (∼5.5 years) given 1) month degree was conferred is not recorded, only year, and 2) thesis defense date was not recorded.

Evaluation of the added resilience course and wellness resources was also limited. The pre- and post-survey for the resilience course was only administered to a single CasePREP Scholar cohort, and the course was only available to two cohorts. While limited in sample size, the data suggest the course was generally well-received and valued. The Scholars appreciated the opportunity to spend time together and with a near peer to discuss non-academic topics important to them at that point in their lives. Feedback from the group exit interview identified tangible targets for course revision or development should CasePREP have continued. As important, feedback also revealed a more general need for social support outside of class. These latter data are in agreement with research that indicates social networks, such as those comprised of friends and peers, are important in persisting in STEM and identifying as a scientist [8].

Despite these limitations, the overall trend is very clear. As one of the longest-running NIGMS-supported PREPs, CasePREP provides a large single-institution dataset available for post-baccalaureate program evaluations. The CasePREP data demonstrate that the majority of Scholars pursued and attained terminal degrees necessary for careers in biomedical research. The data suggest that the US federal government and institutional investments made in these Scholars are economically and scientifically beneficial for multiple stakeholders. Relatively few CasePREP alumni have been lost to follow-up, testament to their persistence in careers in research and the utility of social media in tracking long-term outcomes of trainees [9].

CasePREP, while highly successful, is not unique among NIGMS-supported PREPs. CasePREP is one of more than 60 unique PREPs supported by NIGMS. Published program evaluations demonstrate that Scholars enter PhD or MD/PhD programs at high rates [from “well over half” [2] to “about 65%” [1] to 68% [10] to 73% [11] to “more than 80%” [12] to 84.8% [13] to 86% [14] to 91%) [14,15]] and that their attrition in these PhD programs is relatively low [4.9% [15] to 11% [13] to 23% [2]]. CasePREP outcomes for PhD or MD/PhD program matriculation (83%) and attrition (15.91%) fall within the range published by other PREPs.

The almost uniform success of these NIGMS-supported PREPs, based on matriculation and attrition outcomes, likely reflects a complex interplay between the design and implementation of this research-intense, one-year training program and the Scholars’ motivations and determination to matriculate into a PhD or MD/PhD program. The latter point, in fact, reveals an important selection bias common across PREPs: the successful PREP Scholar applicant, regardless of previous academic performance or research experiences, can articulate or express the explicit intention to enter a PhD program after completion of PREP to become a scientist. PREP Scholars are highly motivated to attend graduate school, and for many, this post-baccalaureate program offered the opportunity for the Scholars to realize their academic and research potentials on par with more well-resourced undergraduates at research-intense institutions. For other Scholars, PREP offered an important transition between undergraduate studies and the rigors of independent graduate study. Surveys and interviews indicate that completion of PREP increased Scholars’ confidence and readiness for graduate school and eventual careers in science [12,16], and this confidence and readiness is reflected in the low PhD program attrition rates documented here and elsewhere for PREPs.

Despite the overwhelming success of PREP, NIGMS early expired the PREP Notice of Funding Opportunity (PAR-22-220) on January 30, 2025, and, citing “changes in NIH/HHS priorities,” abruptly terminated the program by email on April 2, 2025. At the time of the termination email, CasePREP had a scored renewal application in the final stages of award notice. CasePREP’s NIGMS support ended June 30, 2025. To date, NIGMS has not published a revised Notice of Funding Opportunity for a post-baccalaureate research education program or equivalent.

The loss of CasePREP has adverse effects on a variety of stakeholders, including CWRU, Cleveland, Ohio, aspiring scientists, PhD programs across the country, and the scientific community at large. The CasePREP renewal, had it been supported at the requested level, would have brought in ∼$2.7 million dollars spent and taxed in the local, state, and federal economies. The five-year renewal would have supported an additional 30 CasePREP Scholars and would have partially supported two faculty and an administrator. The termination of the program removes a tangible, and, for some Scholars, necessary pathway to graduate school. For PhD programs, the absence of PREP may translate into fewer prepared applicants and high attrition for those who matriculate. The high attrition or lower PhD completion rates, in turn, will lower the return of federal funding investment in producing US scientists. For the potential Scholars, loss of PREP reverberates beyond graduate school admissions, impacting real opportunities for upward economic mobility currently realized for CasePREP alumni scientists in industry and senior positions in academia.

Hard to envision but important to quantify is the potential loss of science and scientific discovery associated with loss of future scientists. Similar to recent efforts that describe the potential negative impact of proposed NIH budget reductions [17], we attempted to describe the science that would have been lost had the CasePREP Scholars not trained in science or entered the scientific workforce. Based on the scientific literature co-authored by CasePREP Scholars, the absence of their foundational science would have impacted approximately 240,000 citations in two generations. While these estimates are extreme in the assumption that the absence of the foundational science would directly eliminate downstream science, even a fraction of missed connections between the foundational science and downstream science has a meaningful and negative impact on timely discoveries and translation potential.

More sobering than the loss of citing science is the loss of the scientists themselves. The traditional pathway to a career in science is long in time spent in education and training, and longer yet for trainees whose pathway to science is not linear. NIH has been the main source of institutional- and individual-level support for undergraduate, graduate, and post-doctoral trainees in biomedical research. As of 2025, many of these training and career development grant awards and mechanisms have been terminated or disappeared (**Appendix E**) [18-20]. All of the NIH predoctoral fellowships or supplements awarded to CasePREP alumni, with the exception of one F30, were associated with a diversity grant mechanism now expired or disappeared. To our knowledge, no active, individual-level fellowship among CasePREP alumni has been terminated. One NIH training supplement to an NIH U01 supporting a CasePREP Scholar was terminated. Anecdotally at least two CasePREP alumni have reported their NRSA F31-D submissions administratively withdrawn prior to NIH study section review. Also, based on a recent LinkedIn search, one CasePREP alumna employed by NIH may have lost their job during the February 2025 reduction in force [21,22]. With severe budget cuts realized at NSF [23,24] and changes made to HHMI [25] and other private fellowships, it is not clear what funding mechanisms outside of NIH are or will be available to predoctoral or postdoctoral trainees from backgrounds uncommon in science. In parallel, anticipated budget cuts to NIH [26,27] and other funding issues [28] are threatening individual institution budgets, forcing most to consider smaller cohorts of trainees compared to past cohorts [29]. The perfect storm of unusual funding and political climates, regardless of diversity funding mechanism availability, is sure to produce fewer future US scientists of all backgrounds for years to come.

If NIGMS-supported PREP has been an experiment, the results show it is a huge success. Collectively, NIGMS PREPs have bridged many gaps between research talents from all walks of life and the unequal access to research experiences necessary to enter graduate training programs on the path to a career in biomedical research. Scientific advancements have greatly benefited from having researchers from different backgrounds and perspectives [30]. Losing NIGMS-supported PREPs, and the future scientists they nurtured, will have devastating effects on the scientific enterprise and society as a whole.

## Acknowledgements

We thank long-time CasPREP administrator Malana Bey and external evaluator Dr. Farren B.S. Briggs. We also thank all past and present PREP directors; past CasePREP Principal Investigators and Directors Drs. Allison K. Hall, Mark Chance, Paul N. MacDonald (emeritus), and Diana Ramirez-Bergeron; CWRU Office of Diversity Initiatives & Community Engagement Director (retired) Joseph T. Williams; CWRU Associate Vice President of Student Affairs and Dean of Students Dr. Dean D. Patterson, Jr. (retired); past CWRU clinical counselor Dr. Elisaida Mendez; CWRU School of Medicine Dean Dr. Stanton L. Gerson; past and present CWRU Vice Deans of Education Drs. Paul N. MacDonald (emeritus), Marvin T. Nieman, and Mark Jackson; past and present Directors of CWRU Medical Scientist Training Program Drs. Clifford V. Harding and Alex Y. Huang; past and present Directors of CWRU Biomedical Scientist Training Program Drs. George R. Dubyak and David T. Lodowski; past and present Directors of CWRU Epidemiology and Biostatistics PhD program Drs. Scott M. Williams and Fredrick R. Schumacher; and Population and Quantitative Health Sciences Chair Dr. Jonathan L. Haines. Finally, we would like to thank the past CasePREP mentors, CWRU course directors, CWRU faculty volunteers, CasePREP Scholars, and CasePREP applicants without whom this program would not have been possible. The findings and conclusions presented in this paper are those of the author(s) and do not necessarily reflect the views of the NIH or the U.S. Department of Health and Human Services. The funders had no role in the preparation of this manuscript or the decision to publish.

