## Supplementary Materials for "PREP-aring is worth it: Success of the Case Western Reserve University Postbaccalaureate Research Education Program and its Scholars"

**Appendix A: Overview of CasePREP**

NIGMS-supported PREP is generally a one-year, non-degree granting post-baccalaureate training period designed to prepare Scholars for graduate school and careers in science through professional development, course work, and research. Similar to other PREPs, CasePREP provided Scholars with a core experience in the laboratory or research group as a scientific apprentice equivalent to a first-year graduate student (**Supplementary Table A1**). Scholars were matched with accomplished investigators and their laboratory or research staff to immerse them in a mentored, research-intensive experience. CasePREP Scholars were expected to take a graduate-level course each semester of the program as well as On Being a Professional Scientist: The Responsible Conduct of Research, a graduate-level course required in the Spring semester. Professional development throughout the academic year included developing and delivering oral presentations (e.g., research updates; journal club), developing and submitting conference abstracts, and presenting scientific posters at local and national conferences. The Scholars also developed Individual Development Plans, wrote personal and research statements for graduate school applications, and drafted their curriculum vitae. In the spring semester, Scholars interviewed for graduate schools after a series of mock interviews with CWRU School of Medicine faculty. Scholars also drafted an NIH-style specific aims page and participated in a mock NIH study section (**Supplementary Table A1**). Throughout the program, Scholars received oral and written feedback from the program directors, program administrator, and research mentors.

**Addition of resilience and wellness resources**

The COVID-19 pandemic and associated isolation and stress revealed a general need for additional mental health support. Through a one-year supplement from NIGMS, CasePREP offered the class of 2023-2024 1) access to counseling services through CWRU's University Health and Counseling Services and 2) a new wellness and resilience course. In the Spring 2024 semester, we implemented the new wellness and resilience curriculum designed around the existing National Institute of Health (NIH) Becoming a Resilient Scientist Series, offered by the Office of Intramural Training & Education (OITE) [1]. This series, consisting of five webinars and discussion groups, was originally developed by NIH in 2020 in response to increased demand for these seminars during the COVID-19 pandemic [1]. The series and discussions covered the following topics**:** Introduction to Resilience and Wellness (Unit 1); Exploring our Self-Talk: Cognitive Distortions and Imposter Fears (Unit 2); Self-Advocacy and Assertiveness for Scientists (Unit 3); Developing Feedback Resilience (Unit 4); and Managing Up to Maximize Mentoring Relationships (Unit 5). All webinars, each two hours in length, are available on-demand through YouTube. Also available were accompanying workbooks designed to emphasize video content and to structure the weekly discussions after videos are viewed. CasePREP's version leveraged the Fall 2022 YouTube videos and associated Fall 2023 workbook materials as weekly assignments. The units were split into two, and the resulting 13 weekly small group discussions were led by an early career trained facilitator whose education and training background mirrored many Scholars' backgrounds ("near-peer"). Neither the CasePREP co-directors nor the CasePREP administrator attended these small group discussions, and all discussions were understood by the Scholars and the facilitator to be confidential.

To evaluate the effectiveness and perceived value of the added resilience and wellbeing resources, a pre- and post-course survey was administered via Google Forms to the CasePREP Scholar cohort using a six-item Brief Resilience Scale (BRS) [2] available through the PhenX Toolkit [3] (protocol ID 251401). The post-course survey also included the open-ended question “Do you have any comments on the resilience in science class or general access to resources related to resiliency during PREP?” In addition to the six-item survey, the facilitator held a group exit interview discussion to ask the following questions: Were you satisfied with the content of the NIH recorded series? What did you think of the format of this semester-long course? Will you use what you learned in graduate school and other personal or professional settings? What, if anything, do you plan to use from this course? What are or might be the factors or barriers to applying what you learned? How can this course be improved? What other resources could be made available related to resilience tools needed to study and work in high-knowledge environments? What part of this course was most useful or valuable to you? The facilitator recorded the discussion to transcribe the conversations, redacting identifying information. The recording was deleted after the transcription was completed. Claude Sonnet 4.5 was used to assist with preliminary categorization of text by sentiment, after which themes were reviewed, refined, and interpreted by the authors. From these data, the CasePREP resilience series was offered for the entire 2024-2025 academic year, and discussion groups met every other week instead of weekly.

***CasePREP Resilience Cohort***

The counseling services and resilience course (see Methods and Materials) were made available with the 2023-2024 cohort of six Scholars. The pre- and post-course survey suggests the majority of Scholars had improved or unchanged resiliency for five of the six items on the brief resilience scale (**Supplementary Figure A1**). The post-course survey open-ended question was answered by the majority and was generally positive ("It was a time to take a breather during the day and it helped me feel more relaxed in the afternoon of that day."; "I enjoyed being able to connect with my peers and bond over shared concerns, it made me feel less alone in my struggles."; "Learning about different resources related to resiliency during PREP and having the opportunity to talk about our experiences/struggles in a safe environment was very helpful. I think having this class made me learn a lot about myself and understand how I can better manage different situations in my life in a healthier way."). Six themes emerged from the guided group exit interview: course content, course structure, application of course content in life as a graduate student, support, relevance, and barriers to implementation (**Supplementary Table A2**). The negative feedback and suggestions included comments related to the course itself and general comments related to support outside of class (**Supplementary Table A2**).

**Supplementary Table A1. CasePREP activities and syllabus, by academic term.** CasePREP activities and formal class syllabus shown here are based on the 2024-2025 cohort. In general, CasePREP Scholars were selected from a competitive application process between January and March or until the slots were filled. CasePREP generally began the first week of July and ended mid-to-late May. All Scholars were hired by CWRU as Research Technicians I, and Scholars were matched with research mentors within weeks of hire. Scholars met as a class with the directors on a weekly basis beginning in the summer. The fall and spring weekly meetings were formalized as a graded, one credit course. Grades were based on class participation, assignments outlined in the table, and completion of Scholars Hub, a custom database designed to capture Scholars' academic activities (e.g., departmental seminar attendance; professional development workshop attendance, classes taken and grades), research products (e.g., abstracts and manuscripts), and progress on graduate school applications. Outside the formal coursework, CasePREP Scholars, directors, and the administrator met for social activities. Overall, CasePREP was designed and scheduled to mimic the heavy courseload, research, and other academic expectations (e.g., journal club, conferences, and oral presentations) required of a first-year graduate student.

| Summer | Fall | Spring |
|---|---|---|
| Onboarding as a CWRU Research Technician 1 | Mentored research | Mentored research |
| CasePREP orientation | Resilience course (no credit hours) every other week | Resilience course (no credit hours) every other week |
| Three 2-day rotations | One 3-4 credit graduate-level course | One 3-4 credit graduate-level course |
| Mentor selection | Personal statements | Responsible Conduct of Research course |
| Social activities | Curriculum vitaes | Graduate school interviews |
| Weekly summer journal club given by graduate students, post-doctoral fellows, and professors | Selection of graduate programs for applications | Introduction to the primary scientific literature |
| Selection of fall semester class | Write and submit abstract for a national scientific conference | Making figures for presentations, manuscripts, grants |
| One-on-one meetings with directors | One-on-one meetings with directors | One-on-one meetings with directors |
| Establish or update LinkedIn account | Apply to at least 9 PhD or MD/PhD programs | Formal journal club |
| Mentored research | Present poster at national scientific conference | Introduction to NIH fellowships |
| | Individual development plans | Social activities |
| | Research oral presentations | Write a Specific Aims page |
| | Mock graduate school interviews with CWRU faculty | Mock study section |
| | Social activities | Research oral presentations |
| | | Select graduate program for fall matriculation |
| | | School of Medicine PREP Research Day presentations |
| | | Network with CasePREP alumni to prepare for transition to graduate school |

**Supplementary Table A2. Resilience cohort group exit interview themes by sentiment.** CasePREP 2023-2024 Scholars were interviewed as a group by the resilience course facilitator. The transcript of the interview was used as input into Anthropic's Claude to identify themes by sentiment.

| Theme | Positive Feedback | Negative Feedback | Suggestions |
|---|---|---|---|
| Course content | Informative and useful for transitioning to graduate school | Too broad; not all content was relevant | More tailored content to specific student needs |
| Course Structure | Flexibility of topics, webinars, and discussions worked well | Content not always aligned with current challenges | Use surveys to adjust weekly topics |
| Application in Graduate Life | Provided valuable tools for mentorship, burnout prevention | Hard to implement without more social support | Follow-up on the implementation of learned strategies |
| Support | Supportive classroom environment; encouraged openness | Not all students had access to social support outside of class | Create more formal support networks in future courses |
| Relevance | Connected to personal experiences and real-time challenges | Topics felt too general at times | Personalize course topics based on student preferences |
| Barriers to implementation | Students valued resilience tools and communication strategies | Mental and emotional barriers prevent application | Provide ongoing support after the course |

**Supplementary Figure A1. Brief resilience scale results before and after a 13-week resilience course.** Six Scholars from the 2023-2024 cohort took the brief resilience scale (BRS) at the beginning of the Spring 2024 semester and at the end of the semester. The BRS is a self-administered, 6-item questionnaire designed and validated to assess the ability to bounce back or recover from stress. Respondents rate each of the following items on a Likert-like scale (1-5): I tend to bounce back quickly after hard times; I usually come through difficult times with little trouble; I tend to take a long time to get over set-backs in my life; it does not take me long to recover from a stressful event; It is hard for me to snap back when something bad happens; I have a hard time making it through stressful events. Scholars' post-survey responses were compared to pre-survey responses to categorize responses as improved (black), unchanged (dashed), or worsened (gray) resiliency (x-axis). Data are expressed as percentages (y-axis).

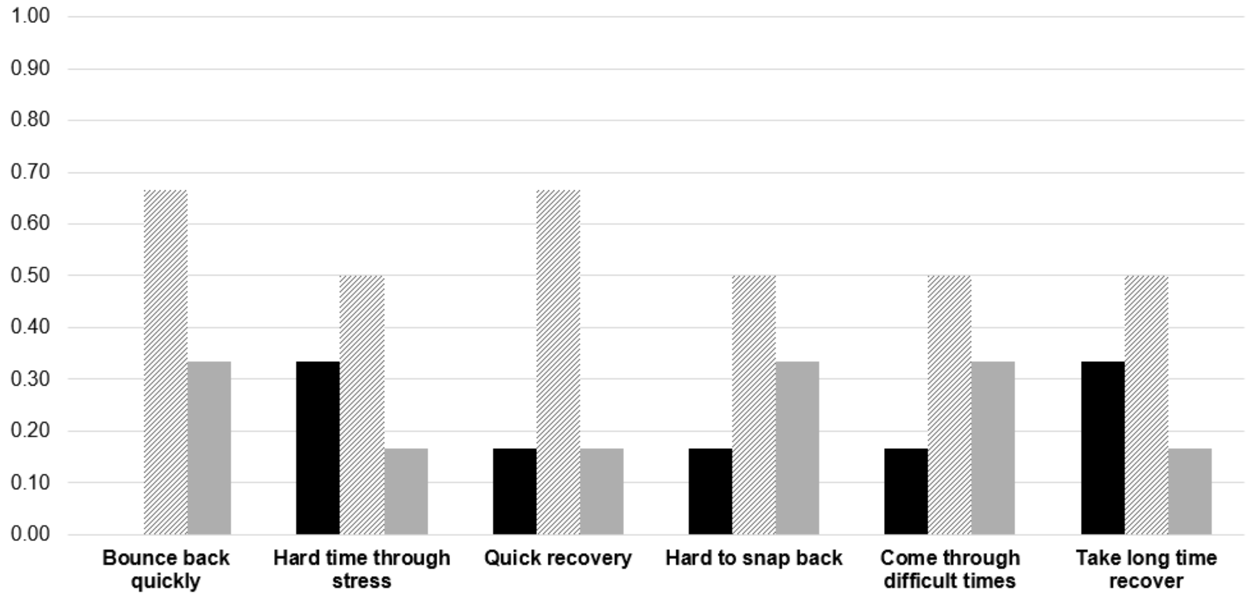

**Appendix B: Economic Impact on the State of Ohio**

Economic impact estimates were based on actual CasePREP salaries committed at the start of each PREP assuming salary payment followed the calendar year (as opposed to the academic year). Adjusted gross incomes were calculated using year-specific federal standard deductions for a single filer for tax years 2007-2024, and federal taxes paid assumed a tax rate of 15%. Historic standard deductions and Social Security and Medicare withholding rates (from irs.gov) were applied to the committed salaries, and Ohio sales tax rates were applied to estimated take home salaries (estimated social security, Medicare, and federal taxes subtracted from committed salaries).

Over the course of 18 years, the NIH awarded a total of $5,623,773 to CWRU to administer CasePREP. Each year of the award, the NIH training grant supported a proportion of the director's (faculty's) effort (15% total 2007-2018; 10% total 2019-2024), a proportion of the program coordinator's (staff's) effort (50%), and the effort for each CasePREP Scholar (100%). In total, the 18-year NIH award supported the equivalent of 119.45 full-time one-year positions at CWRU.

CasePREP was an in-person post-baccalaureate program held on the CWRU campus located in Cleveland, Ohio in Cuyahoga County. The majority of CasePREP Scholars (87.04%) earned their Bachelor's degree from a university located outside of the state of Ohio; consequently, the majority of Scholars became new, tax-paying Ohio residents the year that they participated in CasePREP. Ohio has both a state income tax and a sales tax, the latter of which is applied to apparel, electronics and furniture. The NIH-supported CasePREP salary for Scholars ranged from $21,000 USD (2007-2011) to $27,200 USD (2012-2024). CWRU SOM subsidized CasePREP salaries by $100

USD beginning in 2019, and the subsidy was substantially increased in 2023 ($5,800 USD) and 2024 ($7,800 USD) to closer match national post-baccalaureate salary averages. CasePREP adjustable gross incomes were almost always below the Ohio tax bracket threshold that trigger state tax payment. The sales tax rate for Cleveland, to which all CasePREP Scholars, faculty, and staff were subject, ranged from 7.75% to 8% over the course of CasePREP. Assuming the post-baccalaureate trainees spent 30% of their take home pay on taxable goods, CasePREP Scholars paid an estimated $62,263 in Ohio sales tax over the course of the program. Of the approximately $3,167,600 USD earned in total salary over the 18 years of the program, CasePREP Scholars paid an estimated $189,767 USD and $45,930 USD in Social Security and Medicare withholdings, respectively, and an estimated $319,388 USD in federal taxes. The majority of CasePREP Scholar total salary (~83%), an estimated $2,612,515, was take home pay spent in Ohio on food, housing, sales taxable goods, and other services.

***Return of Investment***

The observed low graduate school attrition rate among CasePREP Scholars (15.91%) suggests early investment in this post-baccalaureate program better ensures that future NIH investment in graduate training results in a trained scientist (e.g., completion of PhD). To quantify this, we first estimated hypothetical NIH funds committed to CasePREP Scholars lost to attrition compared with a hypothetical first-year graduate student cohort of 88 trainees lost to attrition at rates reported in the literature (**Supplementary Figure 1B**). Compared with CasePREP Scholars, NIH investment lost to attrition is 1.57 – 3.46 times higher among typical graduate students, with estimated losses ranging from $2.2 million to $7.26 million USD for a cohort of 88 trainees

(**Supplementary Figure 1B**). Return of investment estimates indicate that NIH costs per CasePREP Scholar who enter graduate school save NIH investments lost to attrition when PhD program attrition rates are >50% (**Supplementary Figure 2B**).

Return of investment was calculated using the standard equation

(Net Program Benefits – Program Costs) / (Program Costs) x 100,

where “program costs” were the NIH per Scholar funds committed to CWRU during the entirely of the program ($5,623,773/108 Scholars = $52,072/Scholar) for the 88 CasePREP Scholars who enrolled in a PhD or MD/PhD program. To calculate “net program benefits”, we defined a benefit of PREP as lower attrition among the 88 Scholars who enrolled in graduate school compared with a hypothetical class of 88 first-year graduate students who did not participate in the post-baccalaureate program. To define the “net program benefits” variable, we assumed graduate students were supported by NIH at $50,000/trainee, which closely represents the total per year stipend, tuition, and research related costs currently allowed for the NIH Predoctoral Individual National Research Service Award (NRSA) and NIH Ruth L. Kirschstein NRSA Institutional Research Training Grant [4]. We estimated NIH support for graduate students who eventually left their programs without a PhD after 1) two years and 2) three years, the periods of training that align with candidacy after comprehensive or qualifying exams, the completion of formal coursework, and the commitment to independent dissertation research. Given attrition rates vary by field of study, program, and year [5], we assumed four attrition rates (25%, 45%, 50%, and 55%) compared with an assumed 16% for CasePREP (15.91% as of March 2026). All economic estimates are conservative and intended to illustrate order-of-magnitude impacts rather than precise accounting.

**Supplementary Figure 1B. Estimated NIH investment in graduate training lost to attrition.** NIH investment, assumed at $50,000/year/graduate student is shown for two (stippled) and three (solid black) years of graduate studies (x-axis) for trainees lost to attrition at varying attrition rates (y-axis). CasePREP attrition rate is the actual rate (15.91%) based on 88 Scholars who entered graduate school as of March 2026. The other attrition rates are applied to a hypothetical cohort of 88 first-year graduate students who did not participate in CasePREP. X-axis represents United States dollars in millions.

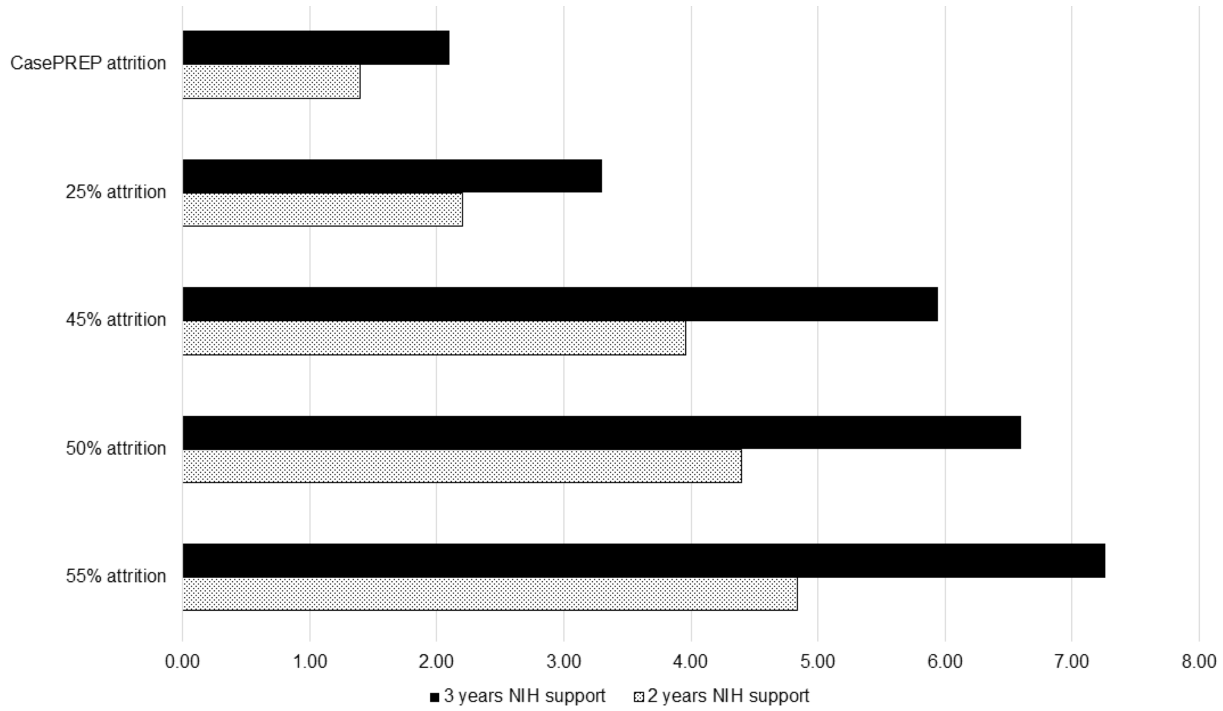

**Supplementary Figure 2B. Return of investment in CasePREP.** Return of investment in CasePREP (y-axis, expressed as %) is estimated, assuming a Scholar attrition rate of 16% compared to various attrition rates for graduate students who did not participate in CasePREP. NIH investment in graduate students is assumed to be $50,000 USD/trainee/year for 1) two years (stippled) and 2) three years (solid black). Net program benefits are estimated as the difference between NIH investment lost to attrition for a hypothetical cohort of 88 graduate students (varying attrition rates) and a hypothetical cohort of 88 CasePREP Scholars (16% attrition). Program costs were estimated as the actual total NIH funds awarded to Case Western Reserve University School of Medicine expressed as a per Scholar cost for the 88 Scholars who entered a PhD or MD/PhD program after CasePREP.

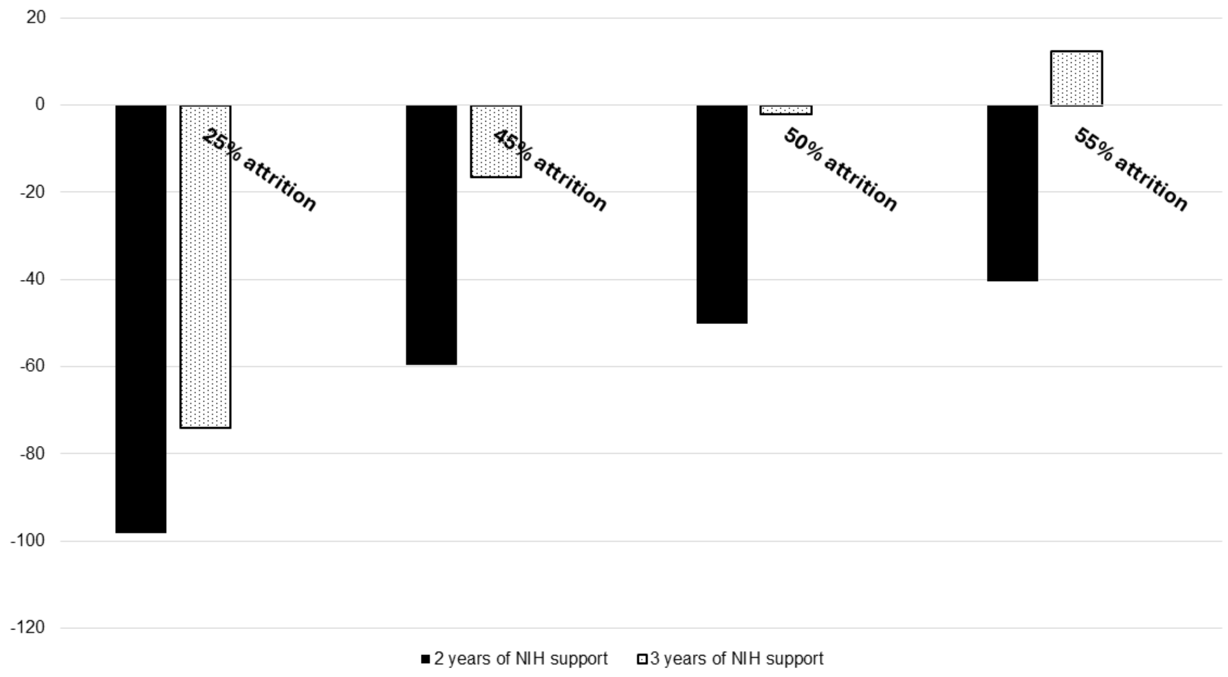

### Appendix C: Institutional Classifications

CasePREP, like all NIGM-supported PREPs at the time, solicited applications from recent college graduates with experiential backgrounds not common in science, including those who were economically disadvantaged. Data indicate that postsecondary education increases wages and promotes intergenerational mobility for those individuals with those credentials or degrees compared with those without postsecondary degrees [6,7], and that the effects are especially pronounced for students from the lowest income quartile [7]. CasePREP Scholars already have a postsecondary education degree (Bachelor's) but are seeking the highest level of education with a doctoral degree. For the CasePREP Scholars who entered a PhD or MD/PhD program, we used the Carnegie Classification of Institutions of Higher Education "student access and earnings" to quantify evidence of social mobility and increased earning potential after completion of PREP.

Carnegie Classifications of Institutions of Higher Education does not provide "student access and earnings" for Schools of Medicine; for these institutions, we used the 2025 US News & World Reports Medical School Research Rankings and assumed all tier 1 medical schools, medical colleges, and Schools of Medicine would be considered "higher earnings". For medical schools not participating in US News & World Reports rankings (e.g., Johns Hopkins University School of Medicine; University of Michigan Medical School; University of Washington), we assumed these would have been classified as tier 1.

Of the 88 CasePREP Scholars who enrolled in a PhD or MD/PhD program, one enrolled in a PhD program at a research institution not included in the Carnegie

Classification of Institutions of Higher Learning and one enrolled in a medical college neither included in the Carnegie Classification of Institutions of Higher Learning nor ranked by US News & World Reports. Of the remaining 86 Scholars, 20 (23.3%) earned a Bachelor's degree with from an institution classified as medium to lower earnings potential and enrolled in a PhD granting program at a university or School of Medicine classified as higher earnings potential.

**Supplementary Figure 1C. Distribution of Institutional Classifications from the Carnegie Classification of Institutions of Higher Education 2025.** Carnegie Classifications of Institutions of Higher Education were used to characterize Scholars' final degree granting undergraduate institutions and institutions enrolled by Scholars for PhD or MD/PhD programs using their "institutional characteristics" and "student access and earnings" columns. According to the Carnegie Classifications of Institutions of Higher Education, the majority of the Scholars' undergraduate institutions are classified as "mixed Undergraduate/Graduate-Doctorate" (43.52%) and "mixed baccalaureate" (17.59%), where "mixed" is defined as <50% of degrees are awarded in any one focus area [8]. Of the Scholars' 58 undergraduate institutions, most were classified as having research activities: "Research 1: Very High Spending and Doctorate Production" (30.56%), "Research 2: High Spending and Doctorate Production" (36.11%); and institutions spending at least 2.5 million on research & development on average in a single year but not classified as Research 1 or 2 (6.48%) [9].

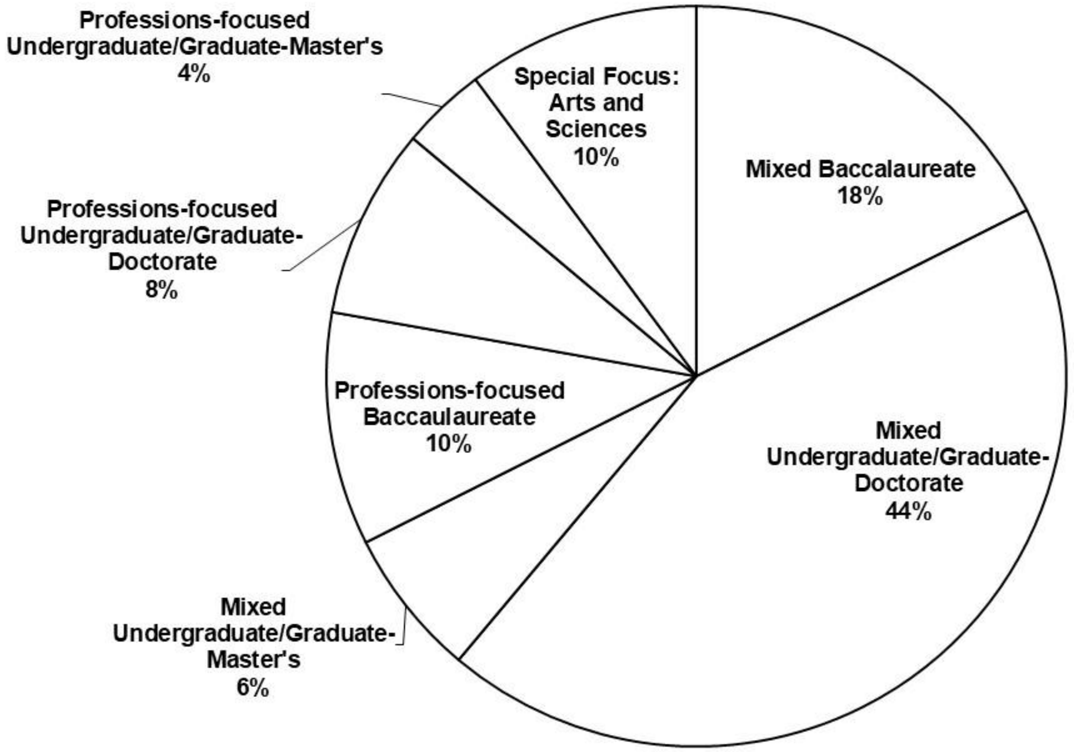
Professions-focused Undergraduate/Graduate-Master's 4%
Special Focus: Arts and Sciences 10%
Mixed Baccalaureate 18%
Professions-focused Undergraduate/Graduate-Doctorate 8%
Professions-focused Baccaulaureate 10%
Mixed Undergraduate/Graduate-Doctorate 44%
Mixed Undergraduate/Graduate-Master's 6%

**Appendix D: CasePREP Scholar research based on publication titles**

PubMed-index publication titles associated with CasePREP Scholars were parsed for 406/407 publications. A word cloud was generated based on the frequency of individual words identified in these titles. One publication was not associated with a parsable title. Publications included here were based on a PubMed search through September 2025.

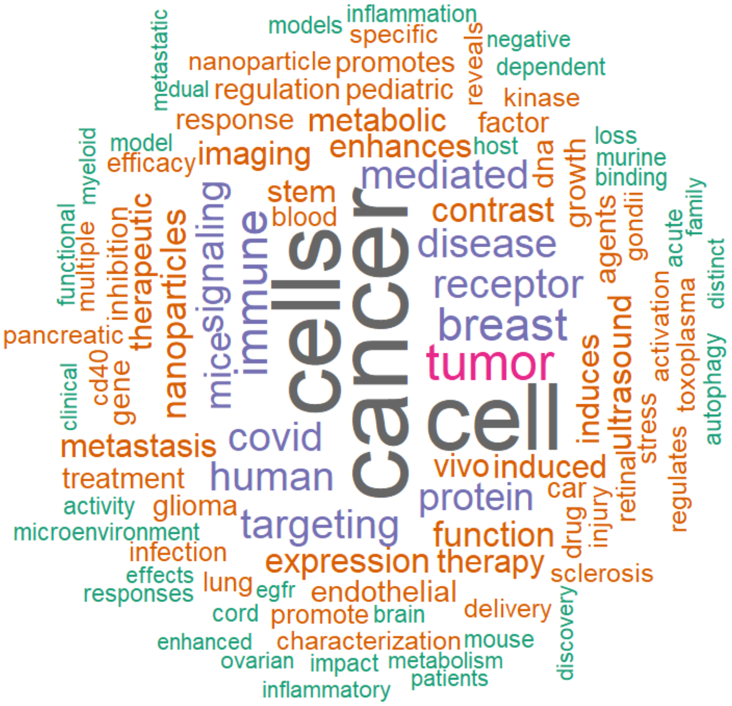

**Appendix E: National Institutes of Health (NIH) Notice of Funding Opportunities (NOFOs) for various training programs**

Shown are NIH NOFOs mentioned in the main text along with their current status. These NIH NOFOs represented funding opportunities for institutional or individual training programs in biomedical research designed to encourage and support trainees from backgrounds not common in science.

| NOFO number | Funding mechanism | Title | Status |
|---|---|---|---|
| PAR-00-139 | R25 | Post-baccalaureate Research Education Program (PREP) | Expired; reissued as PAR-03-140 |
| PAR-03-140 | R25 | NIGMS' Post-baccalaureate Research Education Program | Expired; reissued as PAR-07-432 |
| PAR-17-051 | R25 | Postbaccalaureate Research Education Program (PREP) | Expired; reissued as PAR-20-066 |
| PAR-22-220 | R25 | Postbaccalaureate Research Education Program (PREP) | Early expired and not reissued [10] |
| PAR-24-137 | T34 | Undergraduate Research Training Initiative for Student Enhancement (U-RISE) | Early expired and not reissued [11] |
| PAR-22-125 | T34 | Bridges to the Baccalaureate Training Program (B2B) | Expired and not reissued [12] |
| PAR-24-138 | T34 | Maximizing Access to Research Careers (MARC) | Early expired and not reissued [13] |
| PAR-24-232 | T32 | Bridges to the Doctorate Research Training Program | Early expired and not reissued [14] |
| PAR-24-031 | T32 | Initiative for Maximizing Student Development (IMSD) | Early expired and not reissued [15] |
| PAR-24-032 | T32 | Graduate Research Training Initiative for | Early expired and not reissued [16] |

| | | | |
|---|---|---|---|
| | | Student Enhancement (G-RISE) | |
| PA-23-271 | F31 | Ruth L. Kirschstein NRSA Individual Predoctoral Fellowship to Promote Diversity in Health-Related Research (Parent F31-Diversity) | Early expired and not reissued [17] |
